# Sideroflexins enable mitochondrial transport and metabolism of neutral amino acids

**DOI:** 10.1101/2025.06.18.660357

**Authors:** Samuel Block, Fangtao Chi, Paul C. Rosen, S. Sebastian Pineda, Luise E. Veelken, Alicia M. Darnell, Keene L. Abbott, Izabella A. Pena, Myriam Heiman, Ömer H. Yilmaz, Matthew G. Vander Heiden

## Abstract

Mitochondria contribute to compartmentalized metabolism in eukaryotic cells, supporting key enzymatic reactions for cell function and energy homeostasis. This compartmentalization necessitates regulated metabolite transport across mitochondrial membranes. While many transport proteins have been identified, several mitochondrial transporters remain poorly characterized. Among these are sideroflexins, an evolutionarily conserved family of mitochondrial inner membrane proteins with unclear function. Using CRISPR/Cas9-mediated candidate transporter knockouts coupled with assessment of mitochondrial membrane permeability via a swelling assay, we identify SFXN1, previously implicated in mitochondrial serine transport and iron homeostasis, as an enabler of mitochondrial transport of multiple neutral amino acids, including proline, glycine, threonine, taurine, hypotaurine, β-alanine, and γ-aminobutyric acid (GABA). We further show that SFXN paralogues exhibit substrate-dependent functional overlap, with SFXN2 and SFXN3 partially rescuing loss of SFXN1 function in glycine-related phenotypes, while SFXN2 and SFXN5 partially rescue SFXN1-dependent changes in enabling GABA transport and metabolism. Altogether, these data establish sideroflexins as key regulators of mitochondrial amino acid transport and metabolism.

## Introduction

Mitochondrial amino acid transport is central to human physiology. Aside from supplying amino acids for the synthesis of proteins encoded by the mitochondrial genome, which are essential for respiration and mitochondrial ATP synthesis,^1,2^ mitochondrial amino acid transport influences a wide range of cellular processes that are frequently altered in disease. For example, mitochondrial proline synthesis and export are upregulated in cancer-associated fibroblasts to support collagen synthesis, promoting tumor growth and metastasis^3^. Conversely, impaired mitochondrial proline import and degradation lead to synaptic dysfunction^4^ and are associated with schizophrenia and hyperprolinemia^5^. Non-proteinogenic amino acids also participate in metabolic pathways that span mitochondrial membranes, such as the degradation of the inhibitory neurotransmitter γ-aminobutyric acid (GABA) within glial mitochondria^6^. While several mitochondrial amino acid transporters, primarily those belonging to the SLC25 family, have been identified^7,8^; the proteins that regulate the mitochondrial transport and metabolism of several amino acids remain poorly understood. This includes many neutral amino acids, such as proline, methionine, alanine, cysteine, threonine, asparagine, phenylalanine, tyrosine, and tryptophan, as well as non-proteinogenic amino acids including GABA, taurine, and β-alanine.

Despite data generated nearly 50 years ago suggesting the existence of a mitochondrial carrier with broad specificity for neutral amino acids^9,10^, no carrier has been identified in humans. To investigate how this class of substrates are transported across mitochondrial membranes, we deleted candidate mitochondrial transporters using CRISPR/Cas9 and assessed mitochondrial uptake of proline using a mitochondrial swelling assay. This approach identified SFXN1 as a regulator of mitochondrial permeability to proline, and together with supporting metabolic studies our findings show that sideroflexins enable the mitochondrial transport and metabolism of multiple neutral amino acids including glycine, threonine, taurine, hypotaurine, β-alanine, and GABA.

## Results

### A targeted swelling screen identifies SFXN1 as a regulator of mitochondrial proline permeability

Studies using isolated mitochondria from non-human organisms have provided evidence for the existence of a mitochondrial proline transporter^9,11–13^, but the molecular identity of this transporter and whether a comparable transport mechanism exists in humans remains unknown. To determine whether human mitochondria exhibit proline permeability consistent with transporter-mediated uptake, we isolated mitochondria from HEK293T cells and measured the kinetics of proline uptake using mitochondrial swelling, a well-established assay used historically to assess the properties of many mitochondrial metabolite carriers^14–17^ (Fig. 1A). Compared to sucrose, which does not penetrate the inner mitochondrial membrane, we observed robust swelling with proline (Extended Data Fig. 1A), including a preference for the physiological L isomer (Fig. 1B-C). These data suggest that purified mitochondria with an intact inner membrane possess a proline transport mechanism, since both isomers would elicit equal swelling responses if uptake occurred by diffusion, consistent with studies using mitochondria isolated from rat^9^, yeast^12^ and plants^11^. Furthermore, mitochondria isolated from cells expressing a guide targeting the basic amino acid transporter SLC25A29^7^ demonstrated reduced swelling in response to arginine (Extended Data Fig. 1B), confirming the utility of the swelling assay in assessing mitochondrial amino acid transport.

**Fig. 1.**
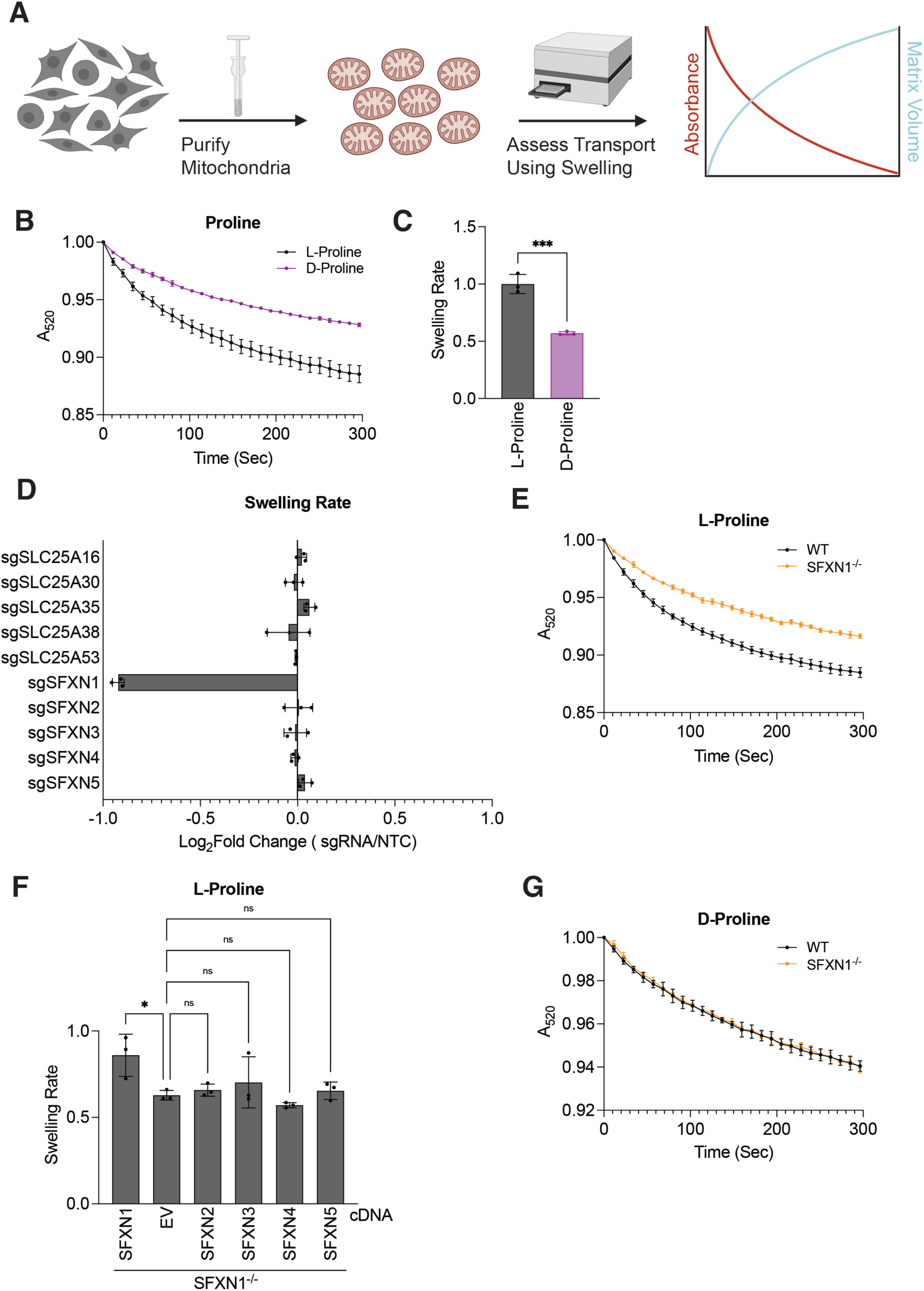
Loss of SFXN1 reduces mitochondrial proline permeability. **(A)** Schematic of mitochondrial swelling experiment to study transport. **(B)** Swelling curves of HEK239T mitochondria treated with either L or D-proline; *n*=3 ± SD. **(C)** Relative swelling rates of HEK239T mitochondria treated with either L or D-proline derived from data shown in (B). Values were normalized to L-proline; *n*=3 ± SD, unpaired t-test \*\*\**p* < 0.001. **(D)** Swelling rates of mitochondria treated with L-proline from HEK293T cells expressing guides targeting the indicated candidate transport protein (sgRNA) compared to non-targeting control (NTC) expressing cells; *n*=3 ± SD. **(E)** Swelling curves of HEK239T mitochondria treated with L-proline from either wild type (WT) or SFXN1 knockout (SFXN1^-/-^) cells; *n*=3 ± SD. **(F)** Swelling rates of mitochondria treated with L-proline from SFXN1 knockout (SFXN1^-/-^) HEK293T cells expressing the indicated cDNA or empty vector (EV); *n*=3 ± SD, one-way ANOVA followed by Dunnett’s multiple comparisons test \**p* < 0.05, ns = not significant. (**G)** Swelling curves of HEK239T mitochondria treated with D-proline from either wild type (WT) or SFXN1 knockout (SFXN1^-/-^) cells; *n*=3 ± SD.

To screen for mitochondrial proline transporter candidates, we used CRISPR/Cas9 gene editing^18^ to individually delete 10 candidate mitochondrial transporters and compared how loss of each candidate affects mitochondrial proline swelling relative to non-targeting control (NTC) (Fig. 1D). In this screen, we considered members of the SFXN protein family (SFXN1-5) and SLC25A38, which have been implicated in the transport of other neutral amino acids^8,19^, as well as SLC25A16, SLC25A30, SLC25A35, and SLC25A53, which are four uncharacterized putative mitochondrial transporters we found to be expressed in HEK293T cells by proteomics (Extended Data Fig. 1C). Of note, mitochondria isolated from cells expressing a sgRNA targeting SFXN1, but not mitochondria isolated from cells expressing a sgRNA targeting any of the other selected transporters, demonstrated reduced swelling with proline compared to NTC (Fig. 1D). To validate that loss of SFXN1 reduces proline swelling, we generated clonal SFXN1 knockout HEK293T and K562 cell lines (Extended Data Fig. 1D-E). Mitochondria isolated from these cells also showed reduced swelling with proline (Fig. 1E, Extended Data Fig. 1F), which was rescued by re-expression of SFXN1 but not overexpression of any other SFXN protein (SFXN2-5) (Fig. 1F, Extended Data Fig. 1G). To assess the stereospecificity of this effect, we also performed the swelling assay with D-proline, which was unaltered by loss of SFXN1 (Fig. 1G). Altogether, these data suggest that mitochondria isolated from human cells exhibit transporter-mediated proline permeability, and that SFXN1 regulates mitochondrial proline permeability.

### Loss of SFXN1 inhibits glutamine-derived proline excretion

Proline can be synthesized in the cytosol through PYCR3 or within mitochondria through PYCR1 or PYCR2^20^, and glutamine is a carbon source for mitochondrial proline synthesis in cultured mammalian cells^21^ (Fig. 2A). However, recent studies have shown that mitochondrial proline synthesis is required to maintain adequate cellular proline levels for proliferation^22,23^. To determine if the reduction in mitochondrial proline permeability we observe in SFXN1 knockout mitochondria is associated with altered glutamine-derived proline excretion, we incubated HEK293T cells with ^13^C_5_-L-glutamine and measured proline released into the media by liquid chromatography-mass spectrometry (LC-MS). For comparison, we also generated cells lacking various combinations of PYCR1-3 (Extended Data Fig. 2A). Loss of SFXN1 markedly reduced M+5 proline released into media to levels similar to those found in PYCR1/2 double knockout cells (Fig. 2B). To test whether loss of SFXN1 reduces labeled proline excretion due to lower labeling of upstream metabolites, independent of its effect on mitochondrial proline release, we assessed labeling of pyrroline-5-carboxylic acid (P5C) from ^13^C_5_-L-glutamine and found no difference in M+5 P5C in SFXN1 knockout cells, even in the absence of PYCR3 (Fig. 2C). Loss of all PYCR enzymes was required to completely inhibit M+5 proline release, whereas loss of PYCR3 in cells lacking SFXN1 had minimal effect (Fig. 2B). These data suggest that PYCR3 is capable of producing proline in the cytosol when PYCR1/2 are absent, and that SFXN1 is required for efficient glutamine-derived proline excretion, but is not essential for mitochondrial proline export, potentially due to the presence of other mechanisms for mitochondrial proline transport. In line with this, cells lacking PYCR1/2, SFXN1, or SFXN1/PYCR3 can proliferate in the absence of proline, whereas PYCR1/2/3 triple knockout cells are proline auxotrophs (Fig. 2D).

**Fig. 2.**
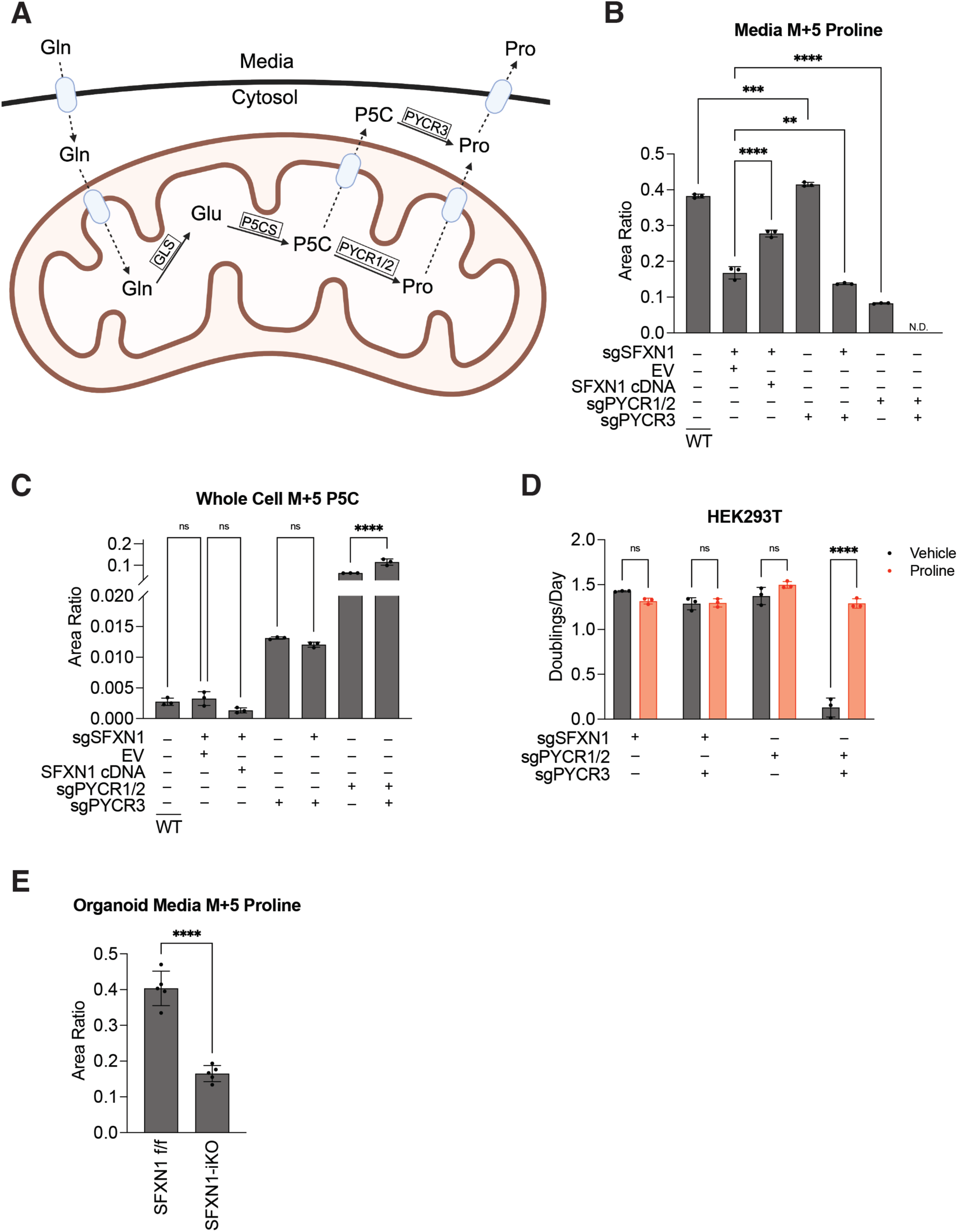
SFXN1 regulates mitochondrial proline export. **(A)** Schematic of how compartmentalized glutamine metabolism relates to proline synthesis. **(B)** Relative media M+5 proline levels measured by mass spectrometry from wild type (WT) HEK293T cells and cells depleted of SFXN1 (sgSFXN1) and the indicated PYCR isoforms (sgPYCR), with or without expression of SFXN1 cDNA or empty vector (EV) as indicated, when cultured in ^13^C_5_-L-glutamine for 24 hours; *n*=3 ± SD, one-way ANOVA followed by Dunnett’s multiple comparisons test \*\**p* < 0.01, \*\*\*\**p* < 0.0001. **(C)** Relative whole cell M+5 P5C levels measured by mass spectrometry from wild type (WT) HEK293T cells and cells depleted of SFXN1 (sgSFXN1) and the indicated PYCR isoforms (sgPYCR), with or without expression of SFXN1 cDNA or empty vector (EV) as indicated, when cultured in ^13^C_5_-L-glutamine for 8 hours; *n*=3 ± SD, one-way ANOVA followed by Šídák’s multiple comparisons test \*\*\*\**p* < 0.0001, ns is not significant. **(D)** Proliferation rates of HEK293T cells depleted of SFXN1 (sgSFXN1) and the indicated PYCR isoforms (sgPYCR) cultured in vehicle or 1mM proline for 48 hours; *n*=3 ± SD, two-way ANOVA followed by Šídák’s multiple comparisons test \*\*\*\**p* < 0.0001. **(E)** Relative media M+5 proline levels measured by mass spectrometry from SFXN1 floxed (f/f) and SFXN1 knockout (-iKO) intestinal crypt organoids^27^ cultured in ^13^C_5_-L-glutamine for 24 hours; *n*=5 ± SD, unpaired t-test \*\*\*\**p* < 0.0001.

Since recent studies reported that sideroflexins regulate the assembly of other mitochondrial protein complexes^24,25^, we considered the possibility that loss of SFXN1 indirectly reduces proline export by altering the levels of other mitochondrial transporters. To test this idea, we used quantitative proteomics to look for changes in protein levels in cells lacking SFXN1, and also cells lacking SFXN4, which has a known role in complex I assembly^25^. While loss of SFXN4 reduced the levels of complex I proteins (Extended Data Fig. 2B-C) as previously described^25^, no significant protein level changes were detected in SFXN1 knockout cells other than SFXN1 itself (Extended Data Fig. 2D).

We next wanted to determine whether SFXN1 regulates proline metabolism in a more physiologically relevant setting. In humans, one source of proline is dietary glutamine, which is metabolized to proline within enterocytes of the small intestine^26^. To study the effects of SFXN1 loss on mitochondrial proline metabolism in this context, we generated SFXN1 knockout intestinal organoids using mice harboring a conditional SFXN1 allele^27^, which was deleted with Cre recombinase expression (Extended Data Fig. 2E). When incubated with ^13^C_5_-L-glutamine, organoids lacking SFXN1 released markedly less M+5 proline (Fig. 2E). Together, these data support a role for SFXN1 in regulating glutamine-derived proline excretion in intestinal cells.

### Neutral amino acids accumulate in SFXN1-null mitochondria

Previous work implicated SFXN1 in mitochondrial serine transport^8^. Given our findings that SFXN1 regulates mitochondrial proline permeability and glutamine-derived proline excretion, we speculated SFXN1 might affect mitochondrial compartmentalization and metabolism of other amino acids. To test this hypothesis, we immunopurified mitochondria^28^ from SFXN1 knockout and control cells (Extended Data Fig. 3A) and measured metabolites using LC-MS. Proline levels were similar in immunopurified mitochondria from control and SFXN1 knockout cells (Extended Data Fig. 3B), possibly due to increased release of upstream metabolites involved in proline synthesis, such as glutamate, which can be exported from mitochondria through multiple known transporters^7^. In support of this, SFXN1 and PYCR1/2 knockout cells excreted increased amounts of M+5 glutamate produced from ^13^C_5_-L-glutamine (Extended Data Fig. 3C). Nevertheless, levels of several neutral amino acids were found to be elevated in SFXN1 knockout mitochondria, including hypotaurine, which is consistent with previous data^29^, as well as elevated asparagine, glycine, β-alanine, taurine and threonine (Fig. 3A).

**Fig. 3.**
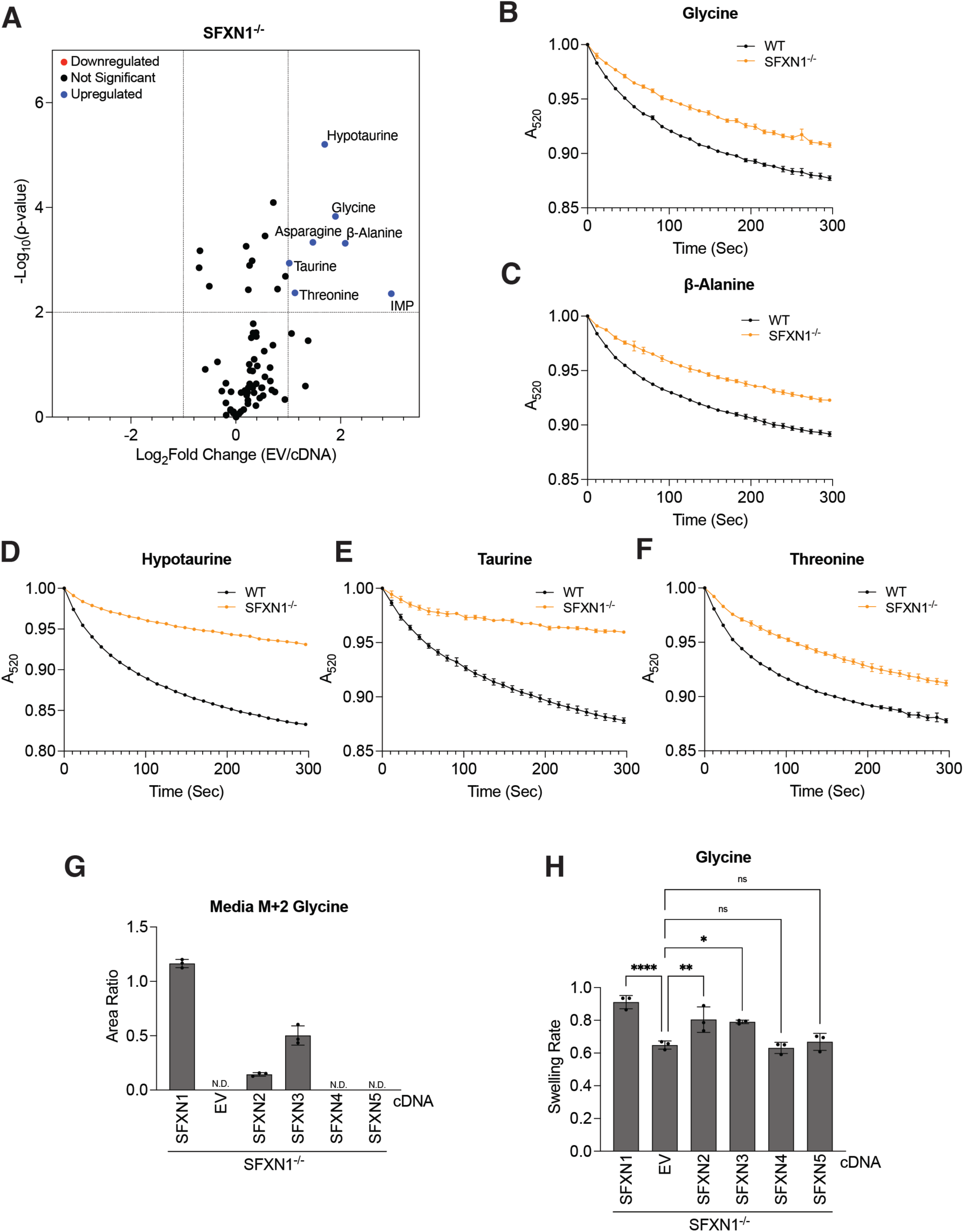
SFXN1 regulates mitochondrial permeability of many polar neutral amino acids. **(A)** Targeted metabolomics of immunopurified mitochondria from SFXN1 knockout (SFXN1^-/-^) HEK293T cells expressing empty vector (EV) compared to SFXN1 knockout cells expressing SFXN1 (cDNA); *n*=3. **(B-F)** Swelling curves of mitochondria treated with the indicated amino acids from wild type (WT) or SFXN1 knockout (SFXN1^-/-^) HEK293T cells; *n*=3 ± SD. **(G)** Relative media M+2 glycine levels measured by mass spectrometry from SFXN1 knockout (SFXN1^-/-^) HEK293T cells expressing the indicated cDNA or empty vector (EV) cultured in ^13^C_3_-L-Serine for 24 hours; *n*=3 ± SD, N.D. is not detected. **(H)** Swelling rates of mitochondria treated with glycine from SFXN1 knockout (SFXN1^-/-^) HEK293T cells expressing the indicated cDNA or empty vector (EV); *n*=3 ± SD, one-way ANOVA followed by Dunnett’s multiple comparisons test \**p* < 0.05, \*\**p* < 0.01, \*\*\*\**p* < 0.0001, ns is not significant.

The buildup of neutral amino acids in SFXN1 knockout mitochondria may reflect a direct or indirect role for SFXN1 in the transport or metabolism of this class of substrates. We directly tested whether SFXN1 affected mitochondrial permeability to these amino acids using mitochondrial swelling assays. Interestingly, mitochondria isolated from SFXN1 knockout cells demonstrated decreased mitochondrial membrane permeability to these amino acids (Fig. 3B-F). However, loss of SFXN1 showed no effect on mitochondrial swelling induced by alanine (Extended Data Fig. 3D), demonstrating amino acid specificity of this permeability phenotype. Since serine undergoes conversion to glycine predominantly within mitochondria via SHMT2^30,31^, and given the reported role of SFXN1 in mitochondrial serine transport^8^, we were surprised that loss of SFXN1 resulted in elevated mitochondrial glycine levels. SFXN1 shares sequence conservation with SFXN2 and 3, and we hypothesized that compensation by other SFXN isoforms, such as SFXN2 expressed by HEK293T cells, might mediate serine transport^8^. In line with this, loss of SFXN1 only mildly reduced mitochondrial swelling in response to serine (Extended Data Fig. 3E), while concomitant loss of SFXN1 and SFXN2 led to a larger decrease in swelling that was rescued by re-expression of SFXN1 (Extended Data Fig. 3F-G). To further probe the effect of SFXN1 in mitochondrial serine/glycine metabolism, we cultured cells with ^13^C_3_-L-serine, which is net consumed^32^ and can be converted into glycine by cytosolic SHMT1 or mitochondrial SHMT2^33^, and measured glycine released into media by LC-MS. Loss of SFXN1 completely inhibited M+2 glycine secretion (Fig. 3G). Over-expression of SFXN2 or SFXN3 was able to partially rescue M+2 glycine secretion in SFXN1 null cells (Fig. 3G), while over-expression of SFXN4 or SFXN5 had no effect, which correlated with their ability to rescue mitochondrial swelling in response to glycine (Fig. 3H)^9^. While it is possible that lower M+2 glycine secretion could be due to lower mitochondrial serine levels, we observed increased levels of serine in SFXN1 knockout immunopurified mitochondria (Extended Data Fig. 3H). It is possible that SFXN1 regulates SHMT2 activity independent of a role in direct serine/glycine transport; however, the SHMT inhibitor SHIN2^34^ completely blocked M+2 glycine production from ^13^C_3_-L-serine (Extended Data Fig. 3I) and had no effect on mitochondrial serine or glycine swelling (Extended Data Fig. 3J-K).

### Sideroflexins regulate mitochondrial GABA permeability and metabolism

Solute carriers often transport multiple chemically similar substrates^35^. Most plasma membrane transporters for proline, glycine, taurine, and β-alanine are members of the SLC6 family, which in humans contains 19 plasma membrane carriers divided among 3 classes: amino acid,monoamine, and GABA transporters^36^. Since the monoamines serotonin, dopamine, and noradrenaline are synthesized in the cytosol^37^, and their degradation occurs on the cytosolic side of the outer mitochondrial membrane^38^, it is unlikely mitochondrial transport of these neurotransmitters is relevant. In contrast, ABAT, the rate-limiting enzyme in GABA degradation localizes to the mitochondrial matrix^39^. Furthermore, the GABA and amino acid transporters within the SLC6 family demonstrate significant overlapping substrate promiscuity^36^, and carrier-dependent GABA transport into mitochondria has been described in rat brain^40^, although the identity of the mammalian mitochondrial GABA transporter has not been reported. To test if SFXN1 is involved in mitochondrial GABA transport, we isolated mitochondria from SFXN1 knockout cells and determined whether GABA-induced swelling was affected. Compared to WT mitochondria, a large reduction in GABA swelling was observed with loss of SFXN1 (Fig. 4A). Importantly, while we observed a slight decrease in membrane potential in the absence of SFXN1 (Extended Data Fig. 4A), GABA swelling was not altered when mitochondria were uncoupled by FCCP treatment (Extended Data Fig. 4B-C), consistent with earlier studies demonstrating the mitochondrial transport of most neutral amino acids is insensitive to membrane potential^10,15,41,42^.

**Fig. 4.**
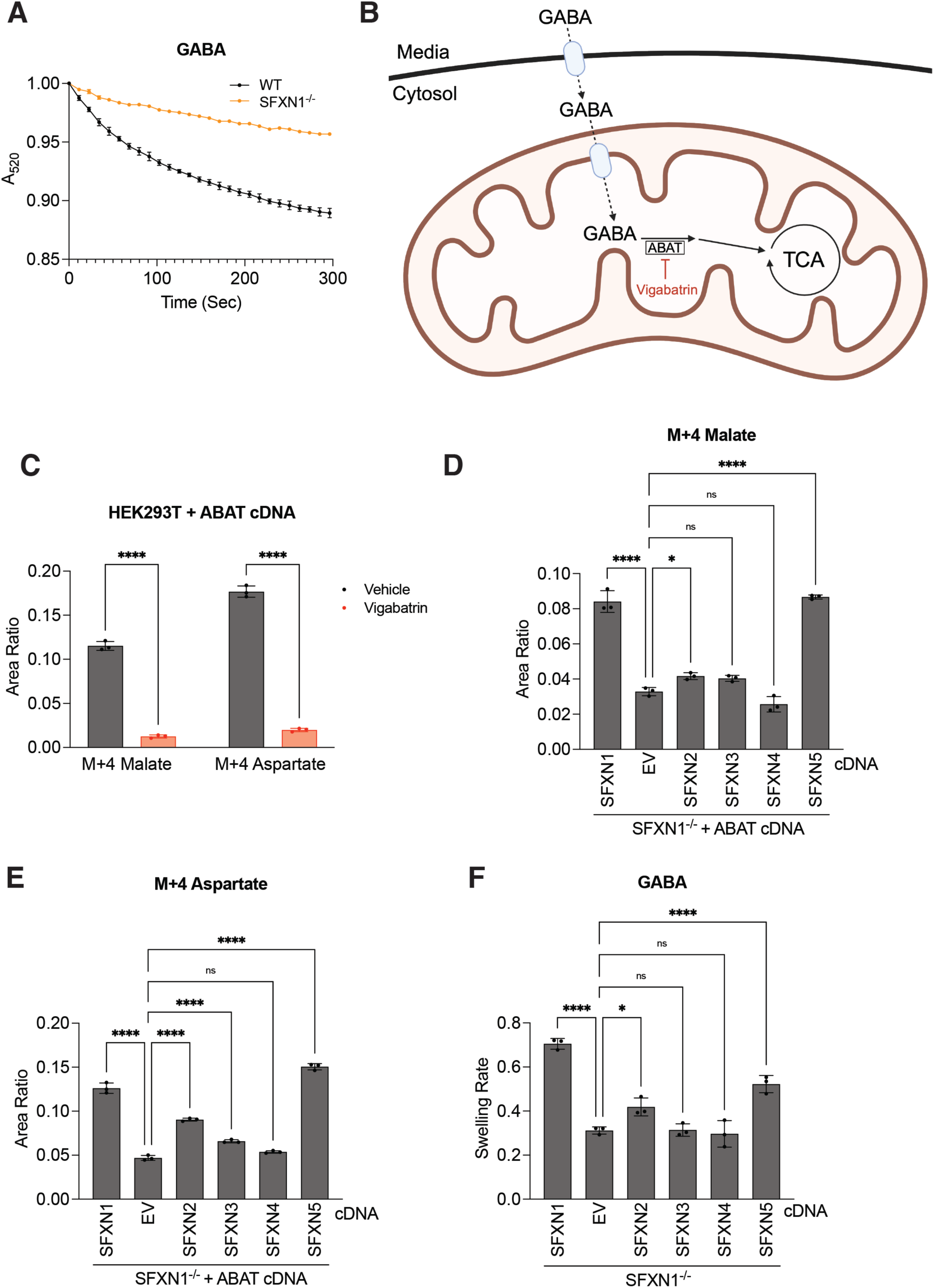
Sideroflexins regulate mitochondrial GABA permeability and metabolism. **(A)** Swelling curves of HEK293T mitochondria treated with GABA from either wild type (WT) or SFXN1 knockout (SFXN1^-/-^) cells; *n*=3 ± SD. **(B)** Schematic of mitochondrial GABA metabolism through the matrix localized enzyme ABAT. TCA is tricarboxylic acid cycle. **(C)** Relative M+4 malate and M+4 aspartate levels measured by mass spectrometry from ABAT cDNA-expressing HEK239T cells that were pre-treated with either vehicle (DMSO) or 1mM Vigabatrin for 24 hours and then cultured in ^13^C_4_-GABA for 30 minutes; *n*=3 ± SD, two-way ANOVA followed by Šídák’s multiple comparisons test \*\*\*\**p* < 0.0001. **(D-E)** Relative whole cell M+4 malate and M+4 aspartate levels measured by mass spectrometry from ABAT cDNA-expressing SFXN1 knockout (SFXN1^-/-^) HEK239T cells expressing the indicated cDNA or empty vector (EV) cultured in ^13^C_4_-GABA for 30 minutes. Area ratios were normalized to ABAT levels determined by western blot densitometry; *n*=3 ± SD, one-way ANOVA followed by Dunnett’s multiple comparisons test \*\*\**p* = 0.001, \*\*\*\**p* < 0.0001, ns is not significant. **(F)** Swelling rates of mitochondria treated with GABA from SFNX1 knockout (SFXN1^-/-^) HEK239T cells expressing the indicated cDNA or empty vector (EV); one-way ANOVA followed by Dunnett’s multiple comparisons test \**p* <0.05, \*\*\*\**p* < 0.0001, ns is not significant.

To examine the effects of SFXN1 loss on GABA metabolism, we expressed ABAT in HEK293T cells (Extended Data Fig. 4D), exposed the cells to ^13^C_4_-GABA and assessed downstream TCA cycle metabolite labeling via the GABA shunt (Fig. 4B), a well-conserved pathway in both eukaryotic and prokaryotic cells^43^. ABAT expression increased M+4 malate and M+4 aspartate levels in HEK293T cells after adding ^13^C_4_-GABA (Extended Data Fig. 4E), which was sensitive to the ABAT-specific inhibitor Vigabatrin^44^ (Fig. 4C). Knockout of SFXN1 inhibited ABAT-dependent labeling from ^13^C_4_-GABA, which was increased by over-expression of SFXN5, and slightly by SFXN2 (Fig. 4D-E), indicating these paralogs may also influence mitochondrial GABA transport. In support of this, over-expression of SFXN2 and SFXN5 also increased GABA swelling in SFXN1 knockout mitochondria, while SFXN3 and SFXN4 had no effect (Fig. 4F).

GABA is the primary inhibitory neurotransmitter in humans and is synthesized in presynaptic neurons and metabolized in glial cells^6^. To determine the expression level and cell type specificity of these SFXN paralogs in GABAergic regions of the brain, we analyzed single nuclear RNA sequencing data obtained from human and mouse striatum^45^, which predominantly contains GABAergic medium spiny neurons^46^, and compared their gene expression profiles to other known GABA metabolic enzymes and transporters. While SFXN1 and SFXN2 are expressed in many cell types within the striatum, SFXN5 expression is largely restricted to astrocytes (Extended Data Fig. 5), which are responsible for recycling synaptic GABA through ABAT^6^. This finding is consistent with previous studies demonstrating SFXN5 is an astrocyte-specific marker^47,48^.

## Discussion

Many enzymatic reactions in eukaryotes are compartmentalized in mitochondria. Consequently, metabolites must be transported across the inner mitochondrial membrane to coordinate metabolism in the mitochondria with reactions in the cytosol. Using mitochondrial swelling and metabolite profiling, we identify the sideroflexins as regulators of mitochondrial transport and metabolism of several neutral amino acids, including proline, glycine, serine, threonine, β-alanine, hypotaurine, taurine, and GABA. While our data suggest that sideroflexins affect the permeability of the mitochondrial inner membrane to these amino acids and are required for their metabolism, we do not provide evidence that they are direct carriers. Interestingly, SFXN4, which has a reported role as a chaperone for complex I mitochondrial proteins^25^, did not rescue any of the permeability defects induced by loss of SFXN1, consistent with its evolutionary divergence from the other sideroflexins^8^. While useful for assessing mitochondrial permeability, swelling assays require supraphysiological substrate concentrations, and future studies such as proteoliposome uptake experiments in reconstituted systems, and investigation of potential protein-protein interactions, are required to determine whether sideroflexins influence transport of these substrates directly or indirectly. However, mitochondrial swelling has been used to assess the activity of specific mitochondrial transport proteins, such as serine transport by SFXN1^49^, as well as Na^+^/H^+^ exchange by the P-module of complex I^50^.

We found that loss of SFXN1 reduces mitochondrial proline permeability and inhibits glutamine-derived proline synthesis in multiple systems. Interestingly, we also found that loss of mitochondrial proline synthesis did not reduce cell proliferation in the absence of exogenous proline, indicating that proliferation-based screens would be ineffective at identifying mitochondrial proline transporters. Mitochondrial proline synthesis has been shown to support cancer cell proliferation by acting as an alternative electron acceptor^21^, and also supports collagen synthesis^51^ and desmoplasia from cancer-associated fibroblasts^3^, which can limit delivery of cancer therapeutics^52^. Interestingly, a co-submitted manuscript indicates that loss of SFXN1-mediated mitochondrial permeability to proline also impairs polyamine biosynthesis and stem cell function in the small intestine^27^. In addition, some cancers may be sensitive to dietary proline restriction^53^. Upregulation of SFXN1 has been found in breast^54^ and lung^55^ cancer, and this is associated with poor prognosis^55,56^. These findings may be due, in part, to SFXN1’s role in mitochondrial proline permeability, although a disruption of other mitochondrial metabolic pathways might also contribute to this phenotype. For instance, loss of SFXN1 also impairs mitochondrial serine-to-glycine conversion, leading to defects in one-carbon metabolism and nucleotide synthesis^8^. Furthermore, we have demonstrated that loss of SFXN1 lowers mitochondrial GABA permeability and inhibits GABA catabolism, and low GABA levels are also predictive of poor prognosis in breast cancer^57^, while the growth of cells isolated from breast cancer-brain metastases are sensitive to Vigabatrin^58^.

Loss of SFXN1 predicts a poor outcome in hepatocellular carcinoma and inhibits toxicity induced by a high-fat diet^59^. This may be due to an inhibitory effect on reactive oxygen species generation^59^, since loss of SFXN1 also protects cells from ferroptosis inducers leading to mitochondrial accumulation of taurine and hypotaurine^29^, which could prevent ferroptosis by acting as antioxidants^29,60–62^. Indeed, apoptosis due to ROS generation by complex I in taurine deficient mice can be rescued with a mitochondrial antioxidant^63^. Our data suggest that hypotaurine and taurine might accumulate in SFXN1 knockout mitochondria due to decreased permeability, providing a potential mechanistic link between SFXN1 loss and ferroptosis resistance, which like apoptosis, can be inhibited by mitochondrial ROS quenching^64^.

We found that SFXN1 loss inhibited isotopically labeled glycine secretion from cells cultured in isotopically labeled serine, in line with previous studies^8,49^. However, we also found that loss of SFXN1 impairs mitochondrial glycine permeability, which could be rescued by SFXN2 and SFXN3. Since SFXN1 was reported as synthetic lethal with loss of cytosolic SHMT1 in media lacking serine^8^, it is possible this phenotype arose from alterations in mitochondrial permeability to serine and glycine, since import of serine and export of glycine is needed for net serine to glycine conversion through mitochondrial SHMT2. Furthermore, loss of function mutations in SFXN1 result in anemia^65^, likely due to impaired heme biosynthesis^24,66^. Since mitochondrial glycine is required for heme biosynthesis^67^, our data suggest that these mutations might impair heme synthesis due to impaired mitochondrial glycine or serine availability. It is nevertheless possible that SFXN1 affects mitochondrial one-carbon metabolism indirectly, and may be context-dependent. Interestingly, loss of function mutations in SLC25A38 also result in anemia^68^, and complete loss of SLC25A38 sensitizes cells to low vitamin B6 which is required for both ALAS^69^ and SHMT^70^ activity, while mitochondrial glycine permeability remains unaffected^49^.

The only mitochondrial GABA transporter that has been characterized is from plants^71^, and while a mutation in aralar was identified that led to an increase in the cytoplasmic/mitochondrial GABA ratio in flies^72^, subsequent data suggested aralar is not a direct GABA transporter^73^. We find loss of SFXN1 results in a mitochondrial GABA permeability defect that impairs GABA degradation, which can be rescued by SFXN5, and partially by SFXN2. SFXN1 and SFXN5 are expressed throughout the brain^8^. SFXN5 expression is largely restricted to astrocytes^47,48^, including in the striatum which contains predominantly GABAergic neurons^46^. Since astrocytes regulate synaptic GABA levels through mitochondrial recycling^6^, drugs targeting SFXN5 could be effective in treating conditions associated with dysregulated GABAergic transmission, such as anxiety disorders and epilepsy, the latter of which is currently treated with the plasma membrane GABA transporter 1 (GAT1) inhibitor Tigabine^74^ and the ABAT inhibitor Vigabatrin^75^.

Furthermore, we also found that striatal astrocytes and neurons in mice and humans express SFXN1 and SFXN2, and neurons may also contribute to GABA recycling^6^.

Lower levels of SFXN1 are found in the brains of Alzheimer’s patients^76^ as well as rats that had undergone ovariectamy^77^, and a P-element insertion in the fly sideroflexin orthologue enhanced Tau-mediated toxicity^76^. Since Alzheimer’s and ovariectomy are associated with dysregulated GABAergic neurotransmission^78,79^, decreased SFXN1 could disrupt GABA homeostasis. However, SFXN1 might also influence these conditions through regulation of other amino acids. Glycine and β-alanine are neurotransmitters, and taurine, proline, and serine have known neuromodulating properties^80^. Since these amino acids are found throughout the body, including GABA^81^, future studies are needed to determine how mitochondrial metabolism influences the levels of these amino acids in different tissues and to what extent the sideroflexins are involved.

## Acknowledgements

We thank all members of the Vander Heiden lab for thoughtful discussions and feedback; Dr. Jonathan Van Vranken and the Thermo Fisher Scientific Center for Multiplexed Proteomics at Harvard Medical School for performing quantitative proteomics and analysis; Dr. Aurora Burds Connor, the KI Preclinical Modeling Core, and the MIT DCM Transgenics Core, including Gong Du, Elmer Umana, and Susan E. Erdman, for generating mouse strains; and Dr. Michael DeMott and the Bioimaging and Chemical Analysis Facility Core at MIT for support with mass spectrometry instrumentation used for polar metabolomics. Figure schematics were created using BioRender.com.

## Author Contributions

S.B., F.C., S.S.P., and L.V. performed experiments and analyzed data. P.C.R., K.L.A. and A.M.D. provided experimental design and critical supplies. Ö.H.Y., M.H. and I.A.P. provided critical feedback. S.B. and M.G.V.H. designed the study and wrote the manuscript with input from all authors.

## Data and materials availability

All data are available in either the main text or supplementary materials. Custom biological materials may be obtained by request from the corresponding author.

## Animal Studies

Mice were under the husbandry care of the Department of Comparative Medicine at the Koch Institute for Integrative Cancer Research at MIT. All procedures were conducted according to AALAC guidelines and approved by the MIT Committee on Animal Care.

## Materials and Methods

### Reagents

Antibodies for SFXN1 (HPA019543), and SFXN2 (HPA026834) were obtained from Atlas Antibodies, FLAG (F1804), and Actin (MAB1501) from Sigma-Aldrich, β-Tubulin (2146), β-Actin (D6A8), Vinculin (E1E9V), and Citrate Synthase (CS) (D7V8B) from Cell Signaling Technology, PYCR3 (OTI1B12) from Novus Biologicals, SFXN1 (12296-1-AP), PYCR1 (13108-1-AP), and PYCR2 (17146) from Proteintech, SFXN4 (PA5-35980) from Invitrogen, and Cytochrome C (Cyt C) (ab13575) from Abcam. Horseradish peroxidase–coupled anti-mouse and anti-rabbit secondary antibody were obtained from Cell Signaling Technologies. For cell culture, DMEM (10-017-CV), RPMI (15-040-CV), and penicillin/streptomycin (30-002-CI) were obtained from Corning, FreeStyle 293 Expression Medium (12338018) was obtained from Gibco, and DMEM without glucose, glutamine, serine, glycine, sodium pyruvate (D9802-01) was obtained from United States Biological. For crypt culture media, Advanced DMEM (12491015), and GlutaMAX (35050061) were obtained from Gibco, EGF (315-09), Noggin (250-38), and R-Spondin (315-32) were obtained from Peprotech, *N*-acetyl-L-cysteine (A9165) and Y-27632 dihydrochloride monohydrate (Y0503) were obtained from Sigma-Aldrich, B27 (17504044) was obtained from Life Technologies, and Chir99021 (C-6556) was obtained from LC Laboratories. For crypt embedding, growth factor reduced Matrigel (356231) was obtained from Corning. Pierce anti-HA Magnetic Beads (88837) were obtained from Thermo Scientific. ^13^C_5_-L-Glutamine and ^13^C_4_-GABA were obtained from Cambridge Isotope Laboratories and all other amino acids, ammonium chloride, FCCP, rotenone, TMRE, potassium EDTA, polybrene, Tris, Sucrose, magnesium chloride, and potassium chloride were obtained from Sigma-Aldrich. Vigabatrin was obtained from Cayman Chemical Company. (Rac)-SHIN2 was obtained from MedChem Express. For transfection, X-tremeGENE 9 was obtained from Roche, Lipofectamine 3000 obtained from Invitrogen, and TransIT-Lenti Transfection Reagent was obtained from Mirus Bio. For LC-MS, HPLC grade water was obtained from Sigma-Aldrich, and HPLC ultra gradient acetonitrile was obtained from Avantor. For cloning, NEB stable competent cells, T4 DNA ligase, BbsI, Esp3I, T4 Polynucleotide Kinase, T4 Ligation Buffer, CutSmart Buffer, and Shrimp Alkaline Phosphatase were obtained from New England Biolabs.

### Cell lines and plasmids

HEK293T and K562 cells were obtained from ATCC. HEK293T cells were used for virus production. ABAT Lentiviral Vector (Human) (CMV) (pLenti-GIII-CMV) (110070610195) was obtained from abm. The plasmids lentiCRISPRv2-Opti (#163126), pSpCas9(BB)-2A-GFP (PX458) (#48138), pU6-(BbsI)_CBh-Cas9-T2A-BFP (#64323), pMXs_FLAG-SFXN1 (#110634), pMXs_FLAG-SFXN2 (#110636), pMXs_FLAG-SFXN3 (#110637), pMXs_FLAG-SFXN4 (#110638), pMXs_FLAG-SFXN5 (#110639), pMXs-bleoR-MitoTag (#218415), pCMV-VSV-G (#8454), pCL-Eco (#12371), pLV hU6-sgRNA hUbC-dCas9-KRAB-T2a-Puro (#71236), pMDLg/pRRE (#12251), pRSV-Rev (#12253), and pMD2.G (#12259) were obtained from Addgene.

### Cell culture

HEK293T cells were passaged in DMEM (10-017-CV) supplemented with 10% fetal bovine serum (FBS), penicillin/streptomycin and Plasmocin® Prophylactic. K562 cells were passaged in RPMI (15-040-CV) supplemented with 10% FBS, 2 mM glutamine and penicillin/streptomycin. For isolating mitochondria for swelling assays, HEK293T cells were grown in suspension in FreeStyle 293 Expression Medium (12338018) supplemented with 1% FBS and penicillin/streptomycin, and K562 cells were grown in suspension in RPMI (15-040-CV) supplemented with 5% FBS, 2mM glutamine, 0.1% Pluronic and penicillin/streptomycin.

### Generation and culture of organoids from Short Conditional intrON (SCON) SFXN1-floxed mice

The SCON SFXN1-floxed mice were generated via microinjection of mice zygotes at the 1-cell stage.^82^ For microinjection, 25 µl of CRISPR injection mix was prepared in nuclease-free buffer (10 mM TRIS-HCl, pH 7.4 and 0.25 mM EDTA), consisting of spCas9 mRNA (100 ng/µl), spCas9 protein (50 ng/µl), sgRNA (50 ng/µl), and ssODN (20 ng/µl, GenScript). The mixture was centrifuged at 13,000 × g at 4 °C for 15–20 min to prevent the clogging of the injection needles. Intestinal organoids were isolated from the resulting mice and cultured in crypt media consisting of Advanced DMEM supplemented with 1x GlutaMAX, 40 ng/ml EGF, 200 ng/ml Noggin, 500 ng/ml R-Spondin, 1 µM *N*-acetyl-L-cysteine, 1x B27, 1 µM Chir99021, and 10 µM Y-27632 dihydrochloride monohydrate as previously described^83^.

### Virus production and transduction

For lentivirus production, HEK293T cells at 80% confluency were transfected with pMDLg/pRRE, pRSV-Rev, pMD2.G, and one of the following: ABAT Lentiviral Vector (Human) (CMV) (pLenti-GIII-CMV), lentiCRISPRv2-Opti, or pLV hU6-sgRNA hUbC-dCas9-KRAB-T2a-Puro with TransIT-Lenti Transfection Reagent. After 48 hours, the virus-containing media was passed through a 0.45 µm filter and stored at −80 °C. For lentiviral transduction, 200,000 HEK293T cells were seeded in 2 mL of culture medium per well in 6-well plates, and after 24 hours the medium was replaced with 1 mL of virus-containing medium. After 24 hours, virus-containing medium was removed and cells expanded into 10 cm plates. 24 hours later, cells were selected with puromycin. Retroviral production and transduction with pMXs plasmids was performed as previously described^49^. Briefly, HEK293T cells were spinfected with retrovirus at 1,200 x g for 45 min at 37 °C. After 18 hours, virus was removed and cells were expanded. Selection medium was added 48 hours post-transduction.

The following oligonucleotides were cloned into lentiCRISPRv2-Opti to generate the cells used in the mitochondrial swelling screen:

sgSLC25A16 (1): 5’-CACCGCGGACCCTAACCATGTCAAG-3’
sgSLC25A16 (2): 5’-AAACCTTGACATGGTTAGGGTCCGC-3’
sgSLC25A30 (1): 5’-CACCGTGCCTGGCGTAACATCGCGG-3’
sgSLC25A30 (2): 5’-AAACCCGCGATGTTACGCCAGGCAC-3’
sgSLC25A35 (1): 5’-CACCGGAGGAACCGACGATAACTCG-3’
sgSLC25A35 (2): 5’-AAACCGAGTTATCGTCGGTTCCTCC-3’
sgSLC25A38 (1): 5’-CACCGGAGGAGCATCTATCACAGTG-3’
sgSLC25A38 (2): 5’-AAACCACTGTGATAGATGCTCCTCC-3’
sgSLC25A53 (1): 5’-CACCGTAAAGTTGGAAACGGCCCCA-3’
sgSLC25A53 (2): 5’-AAACTGGGGCCGTTTCCAACTTTAC-3’
sgSFXN1 (1): 5’-CACCGGTTCCCATTCTCATCCGTGA-3’
sgSFXN1 (2): 5’-AAACTCACGGATGAGAATGGGAACC-3’
sgSFXN2 (1): 5’-CACCGCAGATACAAAGACAGTGCGG-3’
sgSFXN2 (2): 5’-AAACCCGCACTGTCTTTGTATCTGC-3’
sgSFXN3 (1): 5’-CACCGGGGCACAAATCTGCCGACCA-3’
sgSFXN3 (2): 5’-AAACTGGTCGGCAGATTTGTGCCCC-3’
sgSFXN4 (1): 5’-CACCGACGGACTTGATCCCTTTCAG-3’
sgSFXN4 (2): 5’-AAACCTGAAAGGGATCAAGTCCGTC-3’
sgSFXN5 (1): 5’-CACCGCAGTGACAAAGAGTGTGCGA-3’
sgSFXN5 (2): 5’-AAACTCGCACACTCTTTGTCACTGC-3’
sgNTC (1): 5’-CACCGTAACCGATACTCCCCACATT-3’
sgNTC (2): 5’-AAACAATGTGGGGAGTATCGGTTAC-3’

The following oligonucleotides were cloned into pLV hU6-sgRNA hUbC-dCas9-KRAB-T2a-Puro and used with WT HEK293T cells to deplete SLC25A29 and SFXN1^-/-^ HEK239T cells to generate new lines lacking both SFXN1 and either PYCR3 or SFXN2:

sgSLC25A29 (1): 5’-CACCGCGGGCAGGCGCGGTCAGGGA-3’
sgSLC25A29 (2): 5’-AAACTCCCTGACCGCGCCTGCCCGC-3’
sgPYCR3 (1): 5’-CACCGCGGCGTCCGAGGCAACAAGA-3’
sgPYCR3 (2): 5’-AAACTCTTGTTGCCTCGGACGCCGC-3’
sgSFXN2 (1): 5’-CACCGGTAACTGGTGGCATTTGTCC-3’
sgSFXN2 (2): 5’-AAACGGACAAATGCCACCAGTTACC-3’

### Generation of CRISPR knockout cells

Knockout lines were generated using a previously described protocol^84^. Briefly, HEK293T or K562 cells were transfected with pSpCas9(BB)-2A-GFP (PX458) or pU6-(BbsI)_CBh-Cas9-T2A-BFP containing guides of interest. 48 hours later, BFP/GFP positive cells were single cell sorted into 96 well plates and expanded to establish clonal lines. Knockout of genes of interest were confirmed by western blot.

The following oligonucleotides were cloned into either pSpCas9(BB)-2A-GFP (PX458) or pU6-(BbsI)_CBh-Cas9-T2A-BFP to generate clonal knockout lines:

sgSFXN1 (1): 5’-CACCGTCTTCACTGTAACTGACCCC-3’
sgSFXN1 (2): 5’-AAACGGGGTCAGTTACAGTGAAGAC-3’
sgSFXN4 (1): 5’-CACCGGCGCACGTTGGGCTCAATGA-3’
sgSFXN4 (2): 5’-AAACTCATTGAGCCCAACGTGCGCC-3’
sgPYCR1 (1A): 5’-CACCGAGAGCCGGTGCCATCTGTGG-3’
sgPYCR1 (2A): 5’-AAACCCACAGATGGCACCGGCTCTC-3’
sgPYCR1 (1B): 5’-CACCGCTGCTGTGAAGCCCTTGGCC-3’
sgPYCR1 (2B): 5’-AAACGGCCAAGGGCTTCACAGCAGC-3’
sgPYCR2 (1A): 5’-CACCGGATGGTGACACCAGCCGCAC-3’
sgPYCR2 (2A): 5’-AAACGTGCGGCTGGTGTCACCATCC-3’
sgPYCR2 (1B): 5’-CACCGGAACCTGACACGCAGCAACA-3’
sgPYCR2 (2B): 5’-AAACTGTTGCTGCGTGTCAGGTTCC-3’
sgPYCR3 (1A): 5’-CACCGTGGATGGGCGGTGAGCGCAG-3’
sgPYCR3 (2A): 5’-AAACCTGCGCTCACCGCCCATCCAC-3’
sgPYCR3 (1B): 5’-CACCGCCCACGAAGCCCACGCGCCG-3’
sgPYCR3 (2B): 5’-AAACCGGCGCGTGGGCTTCGTGGGC-3’

PYCR1/2 and PYCR 1/2/3 knockout lines were generated using 2 guides per gene

### Immunoblotting

Cell immunoblots were performed as previously described^85^. Briefly, cells were washed 1x in PBS and lysed in RIPA buffer containing protease inhibitors. Proteins were resolved using SDS-PAGE and transferred to nitrocellulose membranes for immunoblotting using the iBlot2 Dry Blotting System (Thermo Fisher). The blots were blocked for 1 hour in TBST with 5% (w/v) non-fat dry milk. Antibodies were also used in TBST with 5% (w/v) non-fat dry milk. Primary antibodies were used at 1:1000 dilution, and HRP-linked secondary antibodies were used at 1:5000 dilution. After imaging, blots were treated with a stripping reagent followed by hydrogen peroxide treatment, and then re-blocked and re-probed for other proteins according to a previously described protocol^86^. Organoid immunoblots were performed using a similar protocol, except blots were blocked in TBST with 5% FBS, SFXN1 was detected using 12296-1-AP (for cell immunoblots HPA019543 was used to detect SFXN1), and proteins were probed on separate blots.

### Mitochondrial isolation and swelling assay

Isolation of mitochondria and the swelling assay were carried out using a previously described protocol^49^. Briefly, 1x 10^9^ cells were washed with cold PBS and resuspended in cold isolation buffer (250 mM sucrose, 10 mM Tris pH 7.5, 10 mM KCl, 1.5 mM MgCl_2_, 1 mM potassium EDTA). Cells were lysed on ice using 30 strokes with a dounce homogenizer and tight fit pestle, and mitochondria were isolated by differential centrifugation. Mitochondria were diluted to 2 mg/mL in room temperature swelling buffer (20 mM Tris pH 7.5, 0.5 mM potassium EDTA, 10 µM rotenone) and incubated at room temperature for 30 minutes before swelling was initiated by adding an equal volume of room temperature swelling buffer containing 500 mM substrate (250 mM final concentration). Swelling was monitored by measuring absorbance at 520 nm using a microplate reader, and swelling rates were determined as the absolute value of the change in absorbance at 520 nm from 0 to 90 seconds. Swelling rates were normalized to simultaneously swollen WT mitochondria unless otherwise stated.

### Assessment of proliferation and TMRE staining

For proliferation experiments, 50,000 HEK293T cells were seeded per well in Poly-D-Lysine coated 6-well plates in 2 mL of DMEM (10-017-CV) supplemented with 10% dialyzed FBS and penicillin/streptomycin. 16 hours later the media was removed and cells were washed 1x with PBS and incubated in fresh media supplemented with either vehicle or 1 mM proline. Separately plated cells were also counted at this time point using a Cellometer Auto T4 Plus Cell Counter (Nexcelcom Bioscience). 48 hours later, cells were counted again and doublings were determined as log_2_ fold change between the counts. For TMRE staining, TMRE fluorescence measurements were performed using a previously described protocol^87^. Briefly, HEK293T cells were incubated in DMEM (10-017-CV) supplemented with 10% dialyzed FBS, penicillin/streptomycin, and 150 nM TMRE for 5 minutes at room temperature. Fluorescence was then measured by flow cytometry using a 561 nm laser.

### Isotope labeled metabolite tracing

For ^13^C_5_-L-glutamine tracing into media metabolite measurements, 200,000 HEK293T cells were seeded per well in Poly-D-Lysine coated 6-well plates in 2 mL of DMEM (17-207-CV) supplemented with 4.5 g/L glucose, 2mM L-glutamine, 10% dialyzed FBS, and penicillin/streptomycin. For ^13^C_5_-L-glutamine tracing into whole cell metabolites, 400,000 cells were used. 16 hours later the media was removed and cells were washed 1x with PBS and incubated in fresh media with 2mM L-glutamine substituted with 2mM ^13^C_5_-L-glutamine. For media, 24 hours after addition of label, 15 µL of media was collected and briefly vortexed with 150 µL of extraction buffer (HPLC grade 80% methanol 20% water containing isotopically labeled amino acid standards). For whole cell metabolite quantification, cells were washed 1x with cold blood bank saline at the indicated time points and lysed with 400 µL of cold extraction buffer and vortexed at 4°C for 10 minutes. Both media and whole cell samples were then centrifuged at 20,000 x g for 10 minutes to remove insoluble material. The supernatant was then filtered through a 10 kDa PES filter and metabolites were quantified by LC-MS. Peak areas were normalized to an isotopically labeled internal amino acid standard to yield area ratios, and these ratios were normalized to cell counts performed using a Cellometer Auto T4 Plus Cell Counter (Nexcelcom Bioscience) on separately plated cells that were treated with identical experimental conditions. ^13^C_3_-L-serine tracing was performed using the same protocol, except cells were cultured in DMEM (D9802-01) supplemented with 4.5 g/L glucose, 3.7 g/L sodium bicarbonate, 2mM L-glutamine, 400 µM L-serine, 10% dialyzed FBS, and penicillin/streptomycin. For tracing media, 400 µM L-Serine was substituted with 400 µM ^13^C_3_-L-Serine. For ^13^C_5_-L-glutamine tracing in organoids, intestinal crypts organoids were isolated and cultured in crypt culture medium for 48 hours, and then placed into fresh media with 1x GlutaMAX substituted with 0.2 mM ^13^C_5_-L-glutamine. After 24 hours 15µl of media was added to 150 µl of extraction buffer which was briefly vortexed and centrifuged at 20,000 x g for 10 minutes. The supernatant was then filtered through a 10 kDa PES filter and metabolites were quantified by LC-MS. Peak areas were normalized to an isotopically labeled internal amino acid standard to yield area ratios, and these ratios were normalized to total organoid protein determined by BCA. For ^13^C_4_-GABA tracing, 400,000 HEK293T cells were seeded per well in Poly-D-Lysine coated 6-well plates in 2 mL of DMEM (10-017-CV) supplemented with 10% dialyzed FBS and penicillin/streptomycin. 16 hours later the media was removed and cells were washed 1x with PBS, and cells were incubated in fresh media supplemented with 1mM GABA. 24 hours later, the cells were washed 1x with PBS and fresh media was added with 1mM GABA substituted with 1mM ^13^C_4_-GABA. Cells were washed 1x with cold blood bank saline at the indicated time points and lysed with 400 µL of cold extraction buffer and vortexed at 4°C for 10 minutes. Samples were then centrifuged at 20,000 x g for 10 minutes to remove insoluble material. The supernatant was then filtered through a 10 kDa PES filter and metabolites were quantified by LC-MS. Peak areas were normalized to an isotopically labeled internal amino acid standard to yield area ratios, and these ratios were normalized to cell counts using separately plated cells that were treated with identical experimental conditions.

### Immunopurification of mitochondria

Mitochondria were immunopurified for metabolomics using a protocol adapted from previous studies^28,49,88^. 10x 10^6^ HEK293T cells were seeded in 150 mm plates in 20 mL of DMEM (10-017-CV) supplemented with 10% FBS and penicillin/streptomycin. 16 hours later the plates were placed on ice, media was removed and cells were scraped into 5 mL of cold KPBS and centrifuged at 250 x g for 3 minutes. All following steps were performed at 4°C. Cell pellets were resuspended in 1 mL of KPBS and lysed with 10 strokes using a syringe attached to a 26G needle and centrifuged at 1100 x g for 2 minutes. 5 µl of supernatant was taken for input immunoblotting, and the rest was rotated with 100 µl of KPBS-equilibrated pierce anti-HA-magnetic beads for 5 minutes. The beads were then bound to a magnet and washed 3x with 1 mL KPBS. Before the third wash was removed, 5 µl was taken for IP immunoblotting. 100 µl of extraction buffer was then added and the beads were vortexed for 10 minutes. The beads were then magnet removed, and excess beads and insoluble material were removed by centrifugation at 20,000 x g for 10 minutes. The supernatant was collected and analyzed by LC-MS. For volcano plot analysis, peak areas were normalized to the summed peak areas of all internal isotopic amino acid standards, which was then normalized to Cytochrome C levels determined by immunoblot densitometry. Statistical analysis for targeted metabolomics was performed using MetaboAnalyst 6.0 operating on one factor default settings^89^. For reporting individual metabolites, peak areas were normalized to an isotopically labeled internal amino acid standard to yield area ratios, and these ratios were normalized to Cytochrome C levels determined by immunoblot densitometry.

### LC–MS metabolite measurements

Polar metabolite profiling was performed using a previously described protocol^90^. Briefly, LC–MS was carried out using a QExactive orbitrap mass spectrometer equipped with a Dionex Ultimate 3000 UPLC system (Thermofisher). All samples were injected onto a SeQuant ZIC-pHILIC 2.1 mm × 150 mm (5 µm particle size) column (Millipore Sigma), and chromatography was performed starting with 20% buffer A (20 mM ammonium carbonate, 0.1% ammonium hydroxide), and 80% acetonitrile. A chromatographic gradient of linearly decreasing acetonitrile concentration was used with the mass spectrometer operating in full scan, polarity-switching mode. Metabolite identification was determined by referencing a library of chemical standards which was constructed using external standard pools^91^, and relative quantification of metabolites was carried out using XCalibur 2.2 software (Thermo Fisher Scientific).

### Quantitative LC/MS proteomics

Mitochondria in isolation buffer or whole cells washed 3x with cold PBS were flash frozen in liquid nitrogen and stored at −80°C. Tandem mass tag MS was carried out by the Thermo Fisher Scientific Center for Multiplexed Proteomics at Harvard Medical School (https://tcmp.hms.harvard.edu). Samples for protein analysis were prepared essentially as previously described^92,93^. Proteomes (from whole cells or isolated mitochondria) were extracted using a buffer containing 200 mM EPPS pH 8.5, 8M urea, 0.1% SDS and protease inhibitors. Following lysis, ∼150 µg of each sample was reduced with 5 mM TCEP. Cysteine residues were alkylated using 10 mM iodoacetimide for 20 minutes at RT in the dark. Excess iodoacetimide was quenched with 10 mM DTT. A buffer exchange was carried out using a modified SP3 protocol^94^. ∼1500 µg of Cytiva SpeedBead Magnetic Carboxylate Modified Particles (65152105050250 and 4515210505250), mixed at a 1:1 ratio, were added to each sample. 100% ethanol was added to each sample to achieve a final ethanol concentration of at least 50%.

Samples were incubated with gentle shaking for 15 mins. Samples were washed three times with 80% ethanol. Protein was eluted from SP3 beads using 200 mM EPPS pH 8.5 containing Lys-C (1:50: Wako, 129-02541) and incubated overnight at room temperature. Samples were further digested with trypsin (1:50; ThermoFisher Scientific, 90305R20) at 37°C with vigorous shaking. Acetonitrile was added to each sample to achieve a final concentration of ∼33%. Each sample was labelled, in the presence of SP3 beads, with ∼360 µg of TMTPro 16-plex reagents (ThermoFisher Scientific). Following confirmation of satisfactory labelling (>97%), excess TMT was quenched by addition of hydroxylamine to a final concentration of 0.3%. The full volume from each sample was pooled and acetonitrile was removed by vacuum centrifugation for 1 hour. The pooled sample was acidified using formic acid and peptides were de-salted using a Sep-Pak 50mg tC18 cartridge (Waters). Peptides were eluted in 70% acetonitrile, 1% formic acid and dried by vacuum centrifugation. The TMT labeled peptides were solubilized in 5% ACN/10 mM ammonium bicarbonate, pH 8.0 and ∼300 µg of TMT labeled peptides were separated by an Agilent 300 Extend C18 column (3.5 μm particles, 4.6 mm ID and 250 mm in length). An Agilent 1260 binary pump coupled with a photodiode array (PDA) detector (Thermo Scientific) was used to separate the peptides. A 45-minute linear gradient from 10% to 40% acetonitrile in 10 mM ammonium bicarbonate pH 8.0 (flow rate of 0.6 mL/min) separated the peptide mixtures into a total of 96 fractions (36 seconds). A total of 96 Fractions were consolidated into 24 samples in a checkerboard fashion and vacuum dried to completion. Each sample was desalted via Stage Tips and re-dissolved in 5% FA/ 5% ACN for LC-MS3 analysis. Proteome data were collected on an Orbitrap Fusion Lumos mass spectrometer (ThermoFisher Scientific) coupled to a Proxeon EASY-nLC 1000 LC pump (ThermoFisher Scientific). Fractionated peptides were separated using a 120 min gradient at 550 nL/min on a 35 cm column (i.d. 100 µm, Accucore, 2.6 µm, 150 Å) packed in-house. MS1 data were collected in the Orbitrap (120,000 resolution; maximum injection time 50 ms; AGC 6 × 10^5^). Charge states between 2 and 6 were required for MS2 analysis, and a 120 s dynamic exclusion window was used. Top 10 MS2 scans were performed in the ion trap with CID fragmentation (isolation window 0.5 Da; Rapid; NCE 35%; maximum injection time 50 ms; AGC 1.5 × 10^4^). An on-line real-time search algorithm (Orbiter) was used to trigger MS3 scans for quantification^95^. MS3 scans were collected in the Orbitrap using a resolution of 50,000, NCE of 55%, maximum injection time of 200 ms, and AGC of 3.0 × 10^5^. The close out was set at two peptides per protein per fraction^95^. Raw files were converted to mzXML, and monoisotopic peaks were re-assigned using Monocle^96^. Searches were performed using the Comet search algorithm against a human database downloaded from Uniprot in May 2021. A 50 ppm precursor ion tolerance, 1.0005 fragment ion tolerance, and 0.4 fragment bin offset for MS2 scans collected in the ion trap. TMTpro on lysine residues and peptide N-termini (+304.2071 Da) and carbamidomethylation of cysteine residues (+57.0215 Da) were set as static modifications, while oxidation of methionine residues (+15.9949 Da) was set as a variable modification. Each run was filtered separately to 1% False Discovery Rate (FDR) on the peptide-spectrum match (PSM) level. Then proteins were filtered to the target 1% FDR level across the entire combined data set. For reporter ion quantification, a 0.003 Da window around the theoretical m/z of each reporter ion was scanned, and the most intense m/z was used. Reporter ion intensities were adjusted to correct for isotopic impurities of the different TMTpro reagents according to manufacturer specifications. Peptides were filtered to include only those with a summed signal-to-noise (SN) ≥ 160 across all TMT channels. The signal-to-noise (S/N) measurements of peptides assigned to each protein were summed (for a given protein). These values were normalized so that the sum of the signal for all proteins in each channel was equivalent thereby accounting for equal protein loading. For volcano plots only proteins with ≥ 3 peptides were reported.

### Statistical analysis

Two tailed t-tests were used for comparing differences between two groups, and Analysis of Variance (ANOVA) followed by either Šídák’s or Dunnett’s multiple comparison tests was used for comparing differences between more than two groups. *p* <0.05 was considered statistically significant, and most analyses and plots were generated using GraphPad PRISM 9.

