## Supplemental Figures and Legends for "Sideroflexins enable mitochondrial transport and metabolism of neutral amino acids"

### Extended Data Figure 1

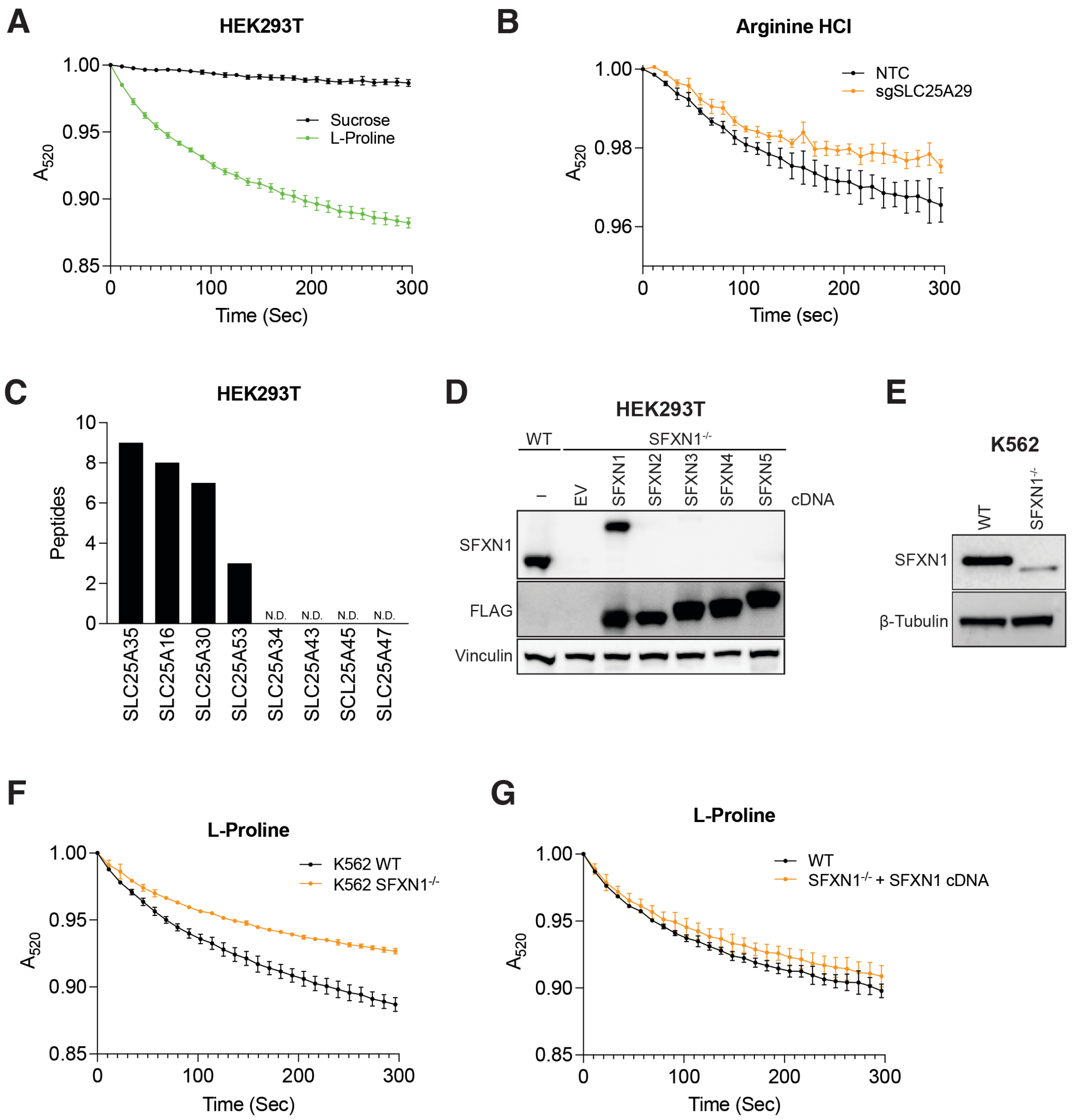

**Extended Data Fig. 1. (A).** Swelling curves of HEK293T mitochondria treated with either sucrose or L-proline; *n*=3 ± SD. **(B)** Swelling curves of HEK293T mitochondria treated with Arginine HCl from either control cells (NTC) or cells expressing a guide targeting SLC25A29 (sgSLC25A29) ; *n*=3 ± SD. **(C)** Peptide counts of the indicated uncharacterized SLC25 family members from HEK293T mitochondria as measured by mass spectrometry; *n*=3, N.D. is not detected. **(D)** Immunoblot of wild type (WT) and SFXN1 knockout ($SFXN1^{-/-}$) HEK293T cells for SFXN1, FLAG and Vinculin as indicated. The $SFXN1^{-/-}$ cells express empty vector (EV) or the indicated SFXN isoform from a cDNA with an N-terminal 3x FLAG-tag. Vinculin was used as a loading control. **(E)** Immunoblot of SFXN1 in wildtype (WT) and SFXN1 knockout ($SFXN1^{-/-}$ ) K562 cells. β-Tubulin was used as a loading control. **(F)** Swelling curves of K562 mitochondria treated with L-proline from either wild type (WT) or SFXN1 knockout ($SFXN1^{-/-}$) cells; *n*=3 ± SD. **(G)** Swelling curves of HEK293T mitochondria treated with L-proline from either wild type (WT) or SFXN1 knockout cells expressing SFXN1 cDNA ($SFXN1^{-/-}$ + SFXN1 cDNA); *n*=3 ± SD.

### Extended Data Figure 2

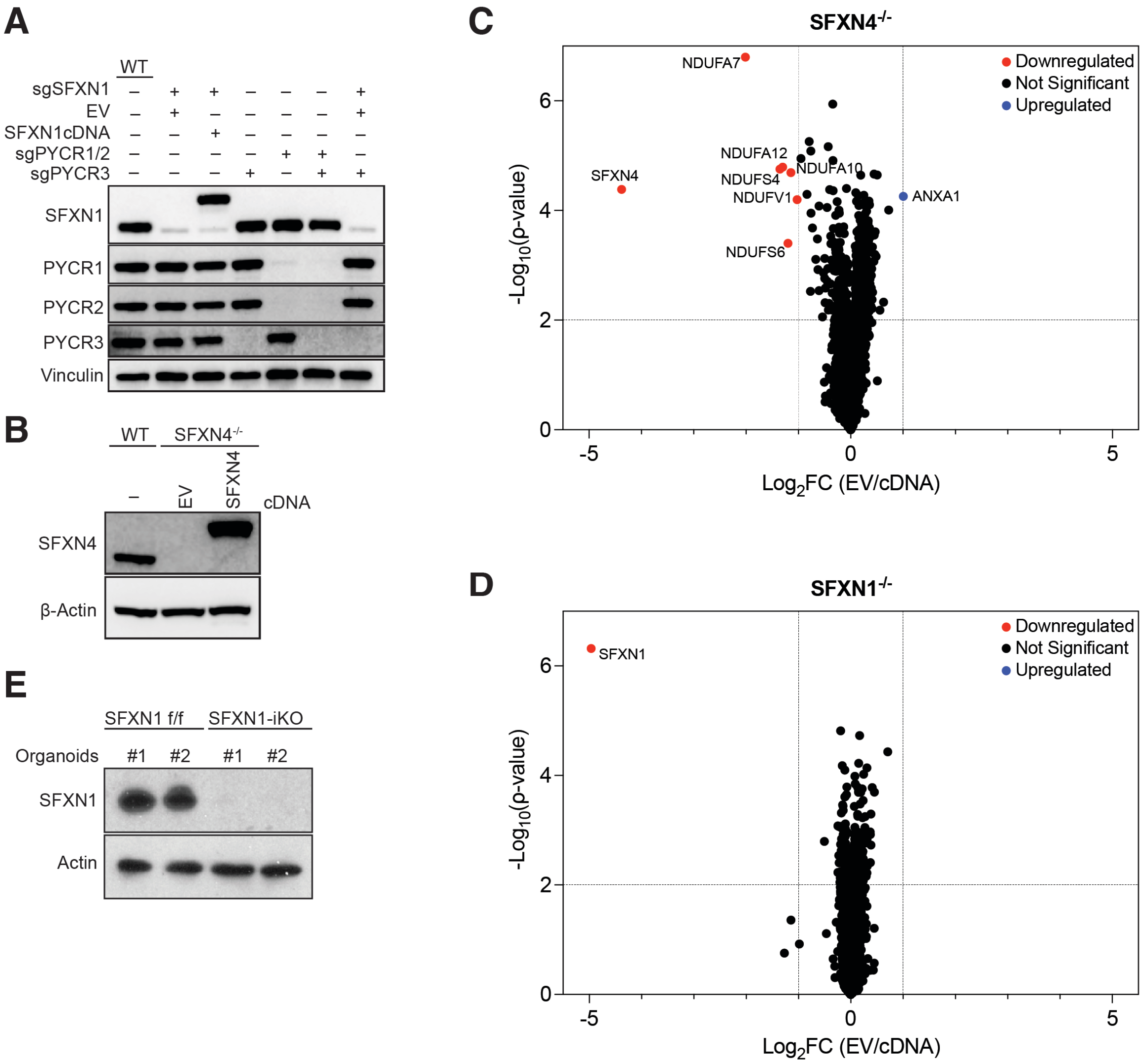

**Extended Data Fig. 2. (A)** Immunoblot for the indicated proteins from wild type (WT) HEK293T cells, or HEK293T cells depleted of SFXN1 (sgSFXN1) and/or the indicated PYCR isoforms (sgPYCR), with or without expression of SFXN1 from a cDNA as specified. EV specifies cells expressing the empty vector control for the cDNA construct. Vinculin was used as a loading control. **(B)** Immunoblot of wild type (WT) or SFXN4 knockout ($SFXN4^{-/-}$) HEK293T cells for the indicated proteins. $SFXN4^{-/-}$ HEK293T cells express either empty vector (EV) or SFXN4 cDNA with an N-terminal 3x-FLAG tag. β-Actin was used as a loading control. **(C)** Quantitative proteomics comparing specified protein levels of SFXN4 knockout ($SFXN4^{-/-}$) HEK293T cells expressing empty vector (EV) to SFXN4 knockout cells expressing SFXN4 (cDNA) as indicated; $n$=3. **(D)** Quantitative proteomics comparing specified protein levels of SFXN1 knockout ($SFXN1^{-/-}$) HEK293T cells expressing empty vector (EV) to SFXN1 knockout cells expressing SFXN1 (cDNA) as indicated; $n$=3. **(E)** Immunoblot for the indicated proteins from two independently derived SFXN1 floxed (f/f) and SFXN1 knockout (-iKO) intestinal crypt organoids[27]. Actin was used as a loading control.

### Extended Data Figure 3

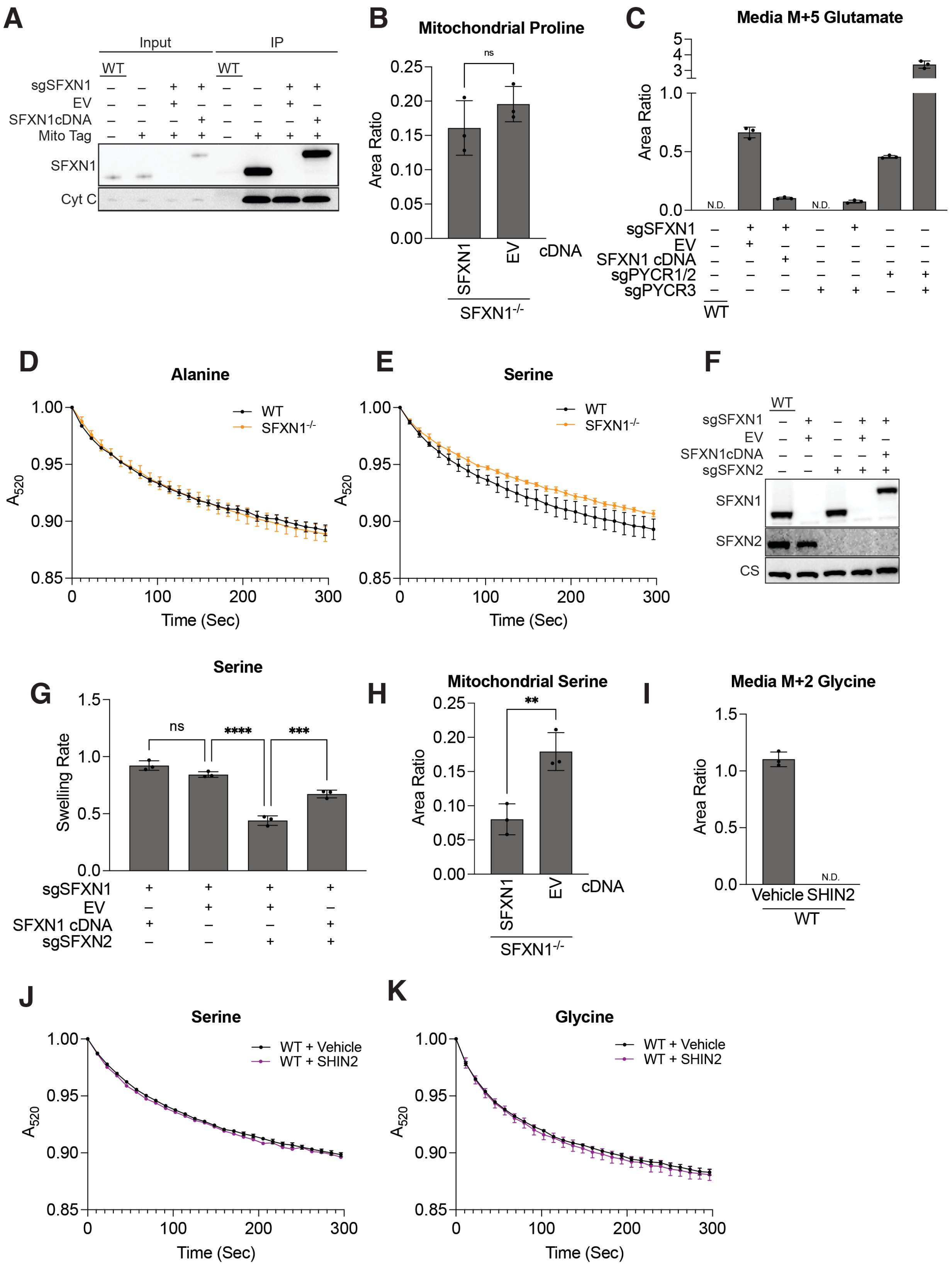

**Extended Data Fig. 3. (A)** Immunoblot for SFXN1 or cytochrome c (Cyt C) of whole cell lysates (Input) and of immunoprecipitated mitochondria (IP) from either wild type (WT) or Mito-tag expressing HEK293T cells, that were or were not depleted of SFXN1 (sgSFXN1), with or without expression of SFXN1 from a cDNA, or empty vector (EV) control as specified. **(B)** Relative proline levels measured by mass spectrometry from immunoprecipitated mitochondria from SFXN1 knockout ($SFXN1^{-/-}$) HEK293T cells expressing either empty vector (EV) or SFXN1 cDNA as indicated; $n$=3 ± SD, unpaired t-test, ns is not significant. **(C)** Relative media M+5 glutamate levels measured by mass spectrometry from wild type (WT) HEK293T cells and cells depleted of SFXN1 (sgSFXN1) and the indicated PYCR isoforms (sgPYCR), with or without expression of SFXN1 from a cDNA or empty vector (EV) as a control, cultured in $^{13}C_5$-L-glutamine for 24 hours; $n$=3 ± SD, N.D. is not detected. **(D)** Swelling curves of HEK293T mitochondria treated with alanine from either wild type (WT) or SFXN1 knockout ($SFXN1^{-/-}$) cells; $n$=3 ± SD. **(E)** Swelling curves of HEK293T mitochondria treated with serine from either wild type (WT) or SFXN1 knockout ($SFXN1^{-/-}$) cells; $n$=3 ± SD **(F)** Immunoblot for the indicated proteins of wild type (WT) and SFXN1 (sgSFXN1) and SFXN2 (sgSFXN2) depleted HEK293T cells expressing the indicated cDNA or empty vector (EV). CS is citrate synthase and was used as a loading control. **(G)** Swelling rates of HEK293T mitochondria treated with serine. Mitochondria were isolated from cells depleted of SFXN1 (sgSFXN1) or SFXN2 (sgSFXN2) and expressing SFXN1 cDNA or empty vector (EV) as indicated; $n$=3 ± SD, one-way ANOVA followed by Šídák's multiple comparisons test ****$p < 0.0001$, ns is not significant. **(H)** Relative serine levels measured by mass spectrometry in mitochondria immunoprecipitated from SFXN1 knockout ($SFXN1^{-/-}$) HEK293T cells expressing SFXN1 cDNA or empty vector (EV) as indicated; $n$=3 ± SD, unpaired t-test **$p < 0.01$. **(I)** Relative media M+2 glycine levels measured by mass spectrometry from wild type (WT) HEK239T cells cultured in $^{13}C_3$-L-serine supplemented either with vehicle (DMSO) or 5 μM (Rac)-SHIN2 for 24 hours; $n$=3 ± SD, N.D. is not detected. **(J-K)** Swelling curves of wild type (WT) HEK293T mitochondria that were pre-incubated with either vehicle (DMSO) or 5 μM (Rac)-SHIN2 for 30 minutes, and then treated with serine or glycine as indicated; $n$=3 ± SD.

### Extended Data Figure 4

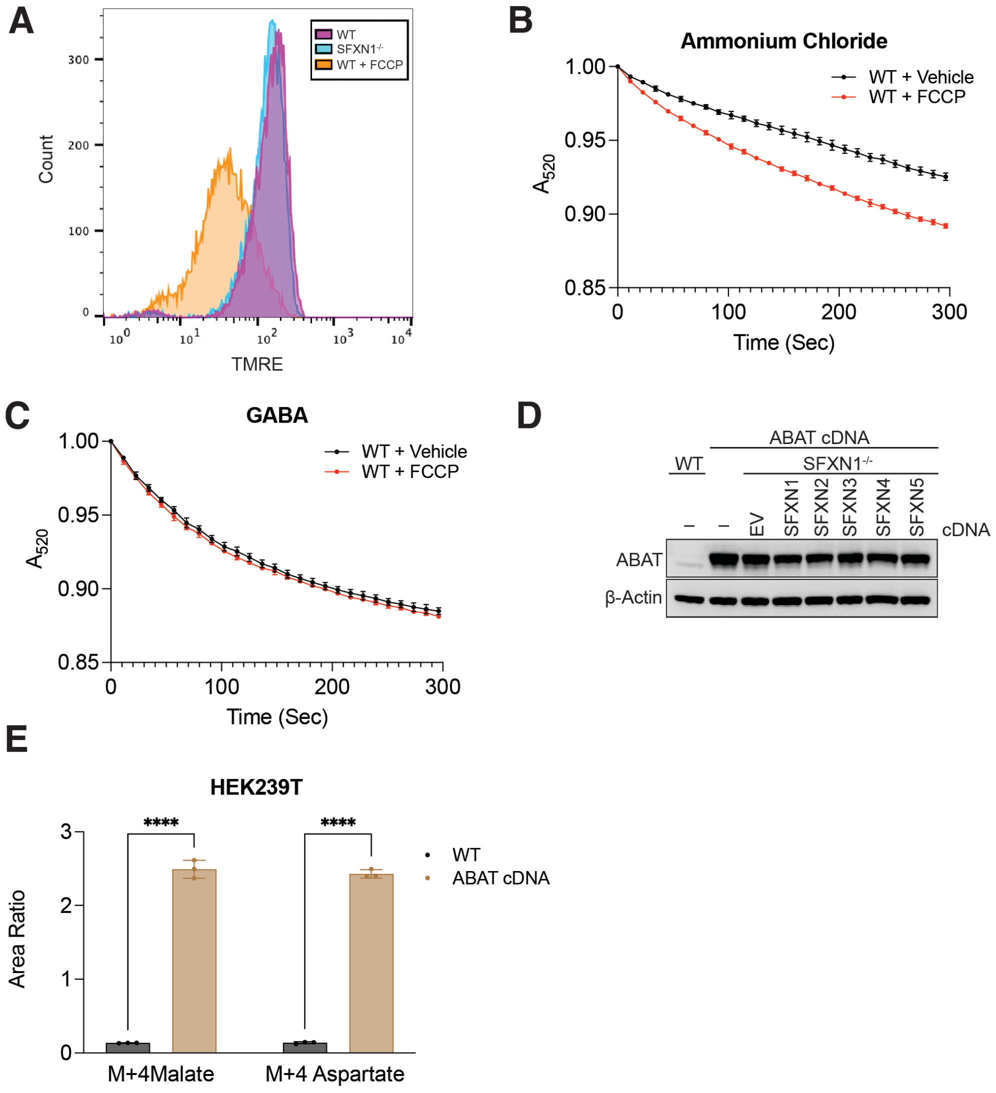

**Extended Data Fig. 4. (A)** Relative mitochondrial membrane potential as measured by TMRE fluorescence using flow cytometry of wild type (WT) or SFXN1 knockout ($SFXN1^{-/-}$) HEK239T cells as indicated. WT cells co-treated with FCCP are also shown as a control; *n*=3. **(B-C)** Swelling curves of wild type (WT) HEK239T mitochondria pre-treated with either vehicle (DMSO) or 20 nM FCCP for 30 minutes and then treated with either ammonium chloride or GABA as indicated; *n*=3 ± SD. **(D)** Immunoblot for the indicated proteins in wild type (WT) or ABAT cDNA-expressing SFXN1 knockout ($SFXN1^{-/-}$) HEK239T cells as indicated. β-Actin was used as a loading control. **(E)** Relative whole cell M+4 malate and M+4 aspartate levels measured by mass spectrometry of wild type (WT) or ABAT cDNA-expressing HEK239T cells cultured in $^{13}C_4$-GABA for 2 hours as indicated; *n*=3 ± SD, two-way ANOVA followed by Šídák's multiple comparisons test ****$p < 0.0001$.

### Extended Data Figure 5

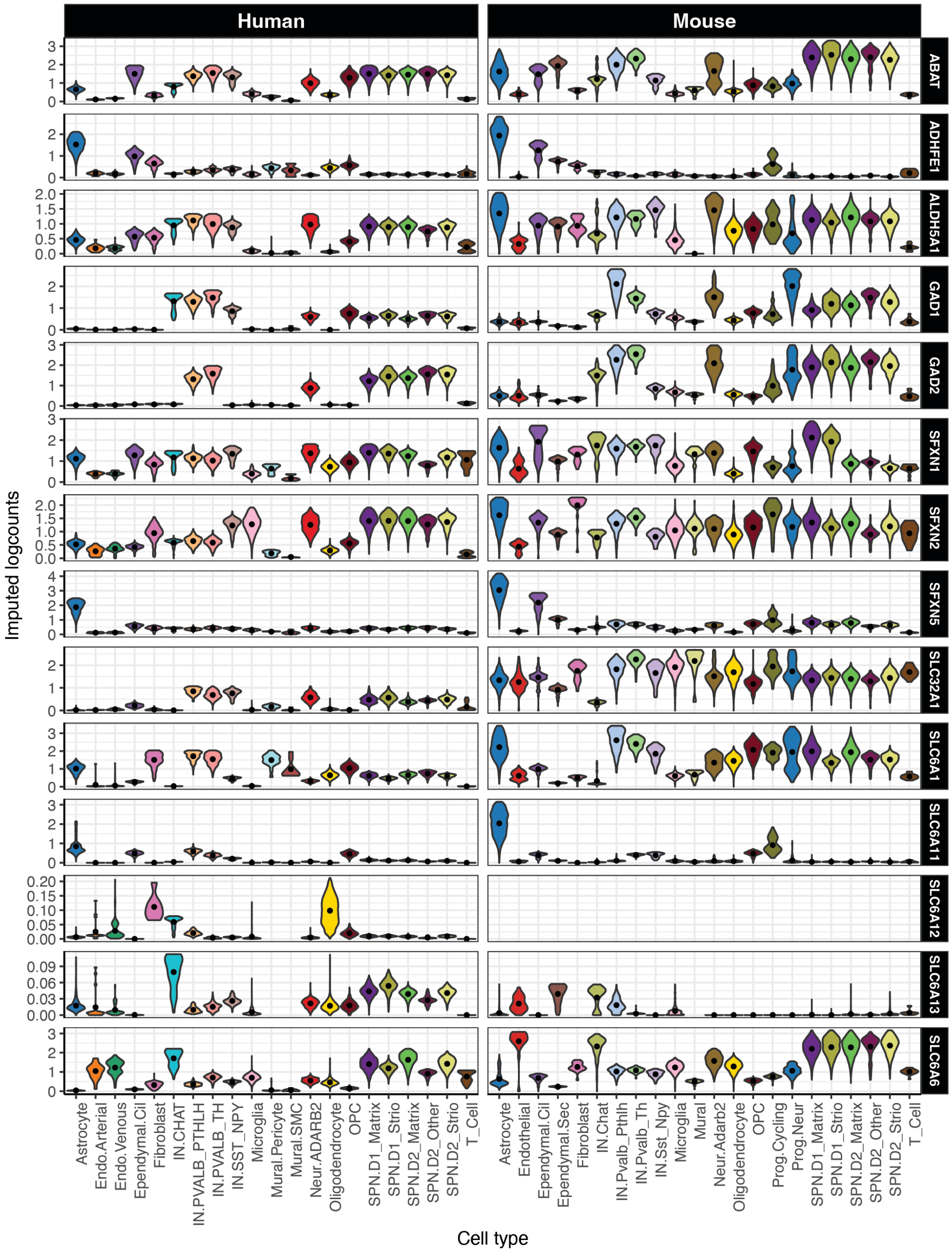

**Extended Data Fig. 5.** Relative levels of the indicated gene transcripts related to GABA metabolism across cell types in human and mouse striatum as derived from published RNA expression data[45].
